# Metabolic regulation of cytokine responses in diffuse large B-cell lymphoma

**DOI:** 10.64898/2026.08.24.746644

**Authors:** Rens Peeters, Aarin White, Sjoerd van Deventer, Annemiek B. van Spriel

## Abstract

Aberrant communication between cells of the immune system can drive disease progression. Cytokines form the central pilar of immune cell communication and are well established factors in lymphomagenesis. An increasing body of evidence suggests that immunometabolism is tightly connected to cytokine production. However, the exact link between metabolism and cytokine responses during lymphomagenesis remains largely unknown. Here, we used established cell models representing the most common form of B-cell lymphoma, diffuse large B-cell lymphoma (DLBCL), to study the effect of metabolism on cytokine production. We found that stimulation or inhibition of the glycolysis pathway could attenuate IL-6, IL-10 and TNFα production by DLBCL. Furthermore, we found that two different subtypes of DLBCL displayed distinct metabolic responses to IL-4. In summary, our work suggests that metabolic pathways could be involved in controlling cytokine production in DLBCL, and paves the road for further research aimed at finding specific metabolic targets that can be exploited for therapeutic intervention.

## Introduction

Communication between cells largely depends on small, soluble glycoproteins and polypeptides called cytokines^1^. Within the immune system, the balance between pro- and anti-inflammatory cytokines affects the outcome of an immune response^2^. Too much pro-inflammatory signalling can result in auto-immunity^3^, whereas too much anti-inflammatory signalling can result in cancer^4^. Aberrant cytokine production can be caused by genetic predispositions^5–7^, epigenetic modifications^8^, environmental factors^9^ and past infections^10^. Moreover, accumulating evidence suggests that cellular metabolism is tightly connected to the ability to generate cytokines^11–13^, as well as respond to them^14,15^.

The modes by which cellular metabolism can affect cytokine profiles are diverse and depend largely on available substrates. The main exogenous substrates for cells are fatty acids (FAs), glucose (Glc), and amino acids (AAs) (Figure 1: block 1, 2 and 3, respectively). Enzymatic processing of these substrates allows cells to generate energy, signalling molecules and precursors for synthesis of nucleotides, proteins and lipids^13,16–18^. Depending on the present needs of a cell, different pathways will be prioritized. For example, intermediates of the glycolysis pathway can either feed into nucleotide or anti-oxidant production via the pentose phosphate pathway (PPP), or into energy production. This process is very rapid and does not require oxygen, but generates only two adenine tri-phosphate (ATP) molecules. Alternatively, substrates can be shuttled into mitochondria, where it fuels the tricarboxylic (TCA) cycle. Intermediates of the TCA cycle are either broken down to efficiently generate energy at the cost of oxygen, or are used for alternative, more anabolic processes (Figure 1, block 4). Interestingly, TCA-intermediates can directly alter the inflammatory phenotype of both innate and adaptive immune cells, by inducing epigenetic modifications in cytokine-associated genes^19,20^. Moreover, nucleotide synthesis pathways depend on TCA-derived amino acids and support cytokine production in infected cells and in immune cells^21,22^. Furthermore, energy (ATP) is required for essential signalling cascades like the protein kinase A (PKA) pathway, and functional lipids are required for both the signalling to-, and the production of cytokines^23,24^. Finally, some metabolic enzymes, like glyceraldehyde-3-phosphate dehydrogenase (GAPDH), were shown to directly inhibit cytokine production by binding to the 3’UTR of mRNA, thereby preventing translation of the cytokine^25^. In conclusion, there is a lot of evidence that links metabolism to the inflammatory phenotype of a cell.

**Figure 1.**
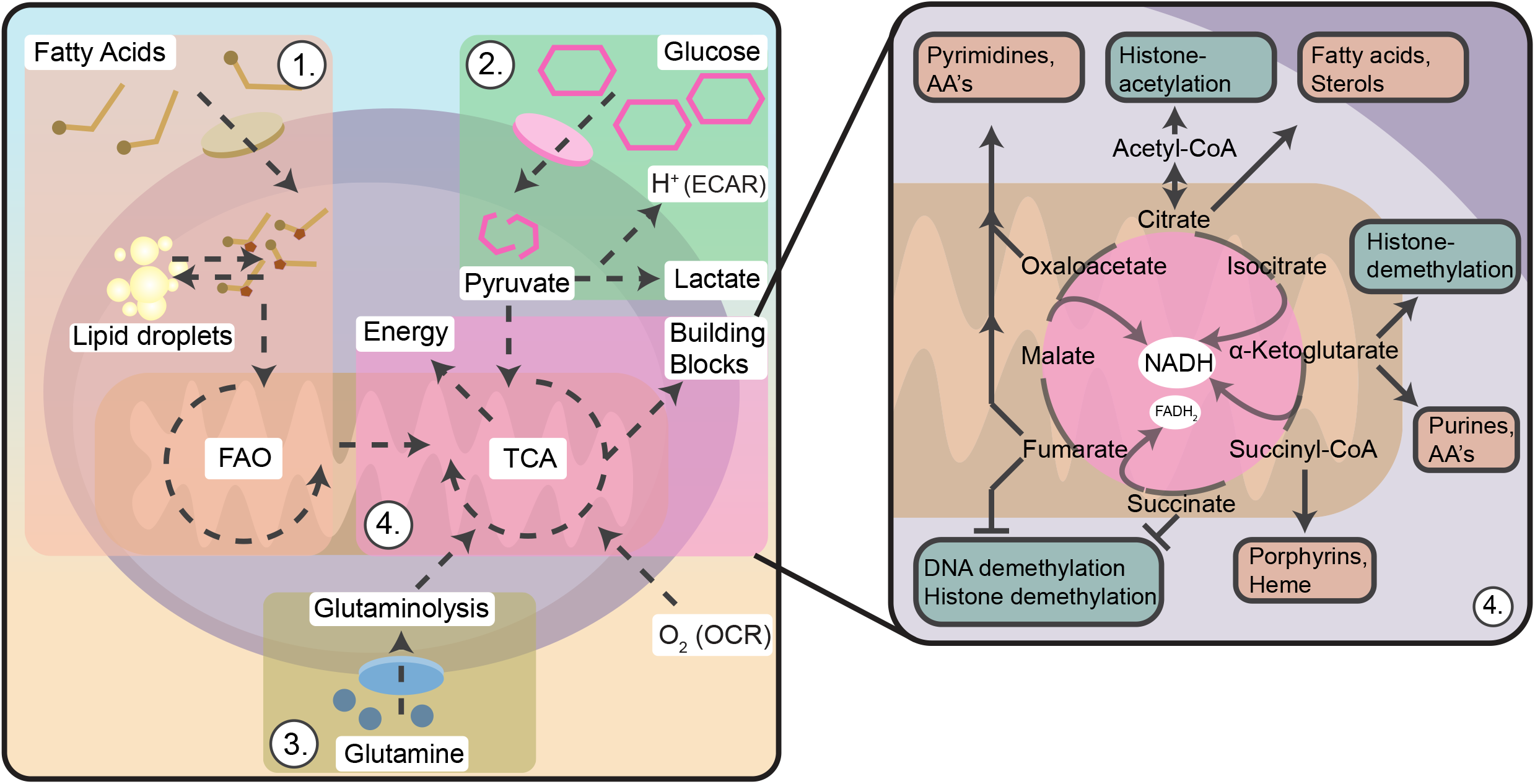
Schematic overview of metabolic pathways and their involvement in cellular processes. Fatty acids (**1)**, glucose (**2)** and glutamine (**3)** are the main substrates used by metabolically active cells. Each substrate is enzymatically processed into either energy or building blocks for proliferation. Glucose via glycolysis, which results in proton secretion. Pyruvate, glutamine and fatty acids can fuel the mitochondria tricarboxylic-acid (TCA) cycle (**4)**. The TCA cycle can generate energy at the cost of oxygen. Alternatively, each intermediate from the TCA cycle can affect cellular processes (**4)**. ECAR: Extracellular acidification rate, OCR: Oxygen consumption rate, CoA: Coenzyme A, AA: Amino acid, NADH: Nicotinamide adenine dinucleotide hydrogen, FADH_2_: Flavine adenine dinucleotide hydrogen.

The rates at which substrates can be taken up and processed is the limiting factor for growth potential. Not surprisingly, cancer cells often display metabolic rewiring in order to sustain their heavy proliferation^26–28^. Aberrant metabolic activity is often observed during lymphomagenesis^29,30^. Interestingly, in patients with the most common form of lymphoma, diffuse-large B-cell lymphoma (DLBCL), increased levels of interleukin-6 (IL-6) are prognostic for inferior disease progression^31,32^. Similarly, elevated levels of interleukin-10 (IL-10) promote tumour growth and immune escape in DLBCL^33^. Furthermore, autoimmune-disease patients treated with inhibitors against tumour-necrosis factor alpha (TNFα), potentially had higher lymphoma incidences as a result of the treatment^34^. These observations illustrate the importance of cytokines in lymphoma, and underscore the need for a better understanding of the processes leading to aberrant cytokine production by these tumours. Although lymphomas often display altered metabolic activity and produce aberrant cytokines, whether and how these two phenomena are related, remains largely unknown.

In the present manuscript, we used established DLBCL cell lines to investigate the potential connection between cancer cell metabolism and cytokine production. We demonstrate that metabolic manipulations like glucose saturation or glycolysis inhibition could alter IL-6, IL-10 and TNFα profiles of DLBCL cell models in vitro. Moreover, we report lymphoma-subtype-specific metabolic changes in response to IL-4. In summary, our results directly correlate cancer-metabolism to cytokine production in DLBCL. Specifically, our results suggest the glycolysis pathway as potential target for therapeutic intervention^35–37^ and invite for more functional research.

## Results

In order to study the effect of cellular metabolism on cytokine production of diffuse large B-cell lymphoma (DLBCL), we treated six different human DLBCL cell lines (BJAB, OciLy8, OciLy19, WSU-NHL, DOHH2 and TMD8) with different metabolic supplements or metabolic inhibitors, alone or in combination with LPS for 24 hours (Figure 2A). As metabolic supplementations cells received saturating amounts of glucose (Glc), glutamine (Gln), palmitate (Palm) or pyruvate (Pyr). To study the involvement of glycolysis, we inhibited the first step of the glycolysis pathway with 2-Deoxy-d-glucose (2-DG). Furthermore, by combining 2-DG with glutamine or pyruvate, we studied whether the glycolysis-intermediates, or rather the end-product, pyruvate, were important for cytokine production. Finally, we assessed the importance of fatty acid (FA) metabolism by inhibiting the FA-activating step of long-chain acyl-CoA synthetase 1 (ACSL1) with TriacsinC (TrcC). After 24 hours, the production of DLBCL-involved cytokines, IL-6, IL-10 and TNFα, was measured by enzyme-linked immunosorbent assays (ELISAs).

**Figure 2.**
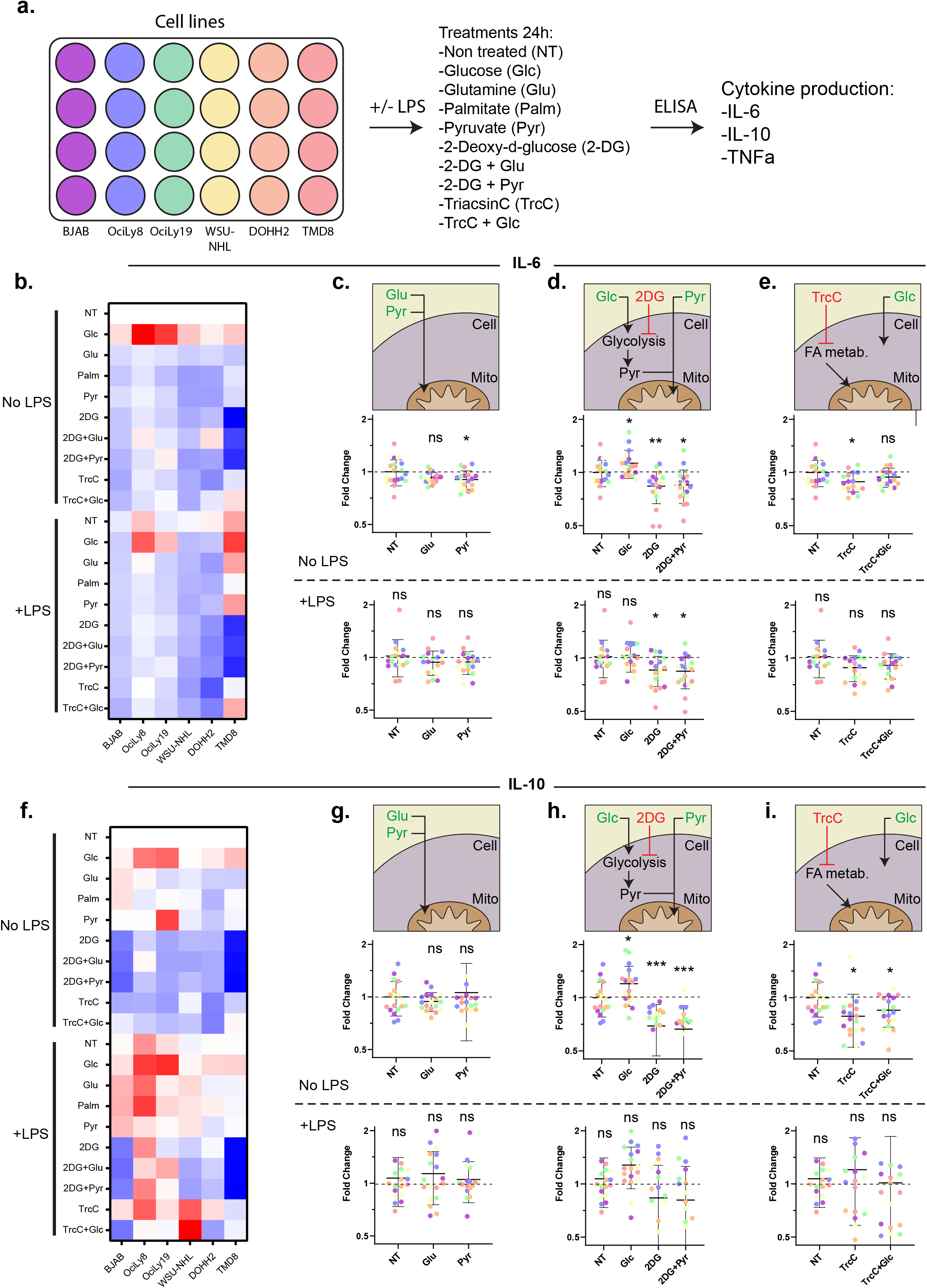
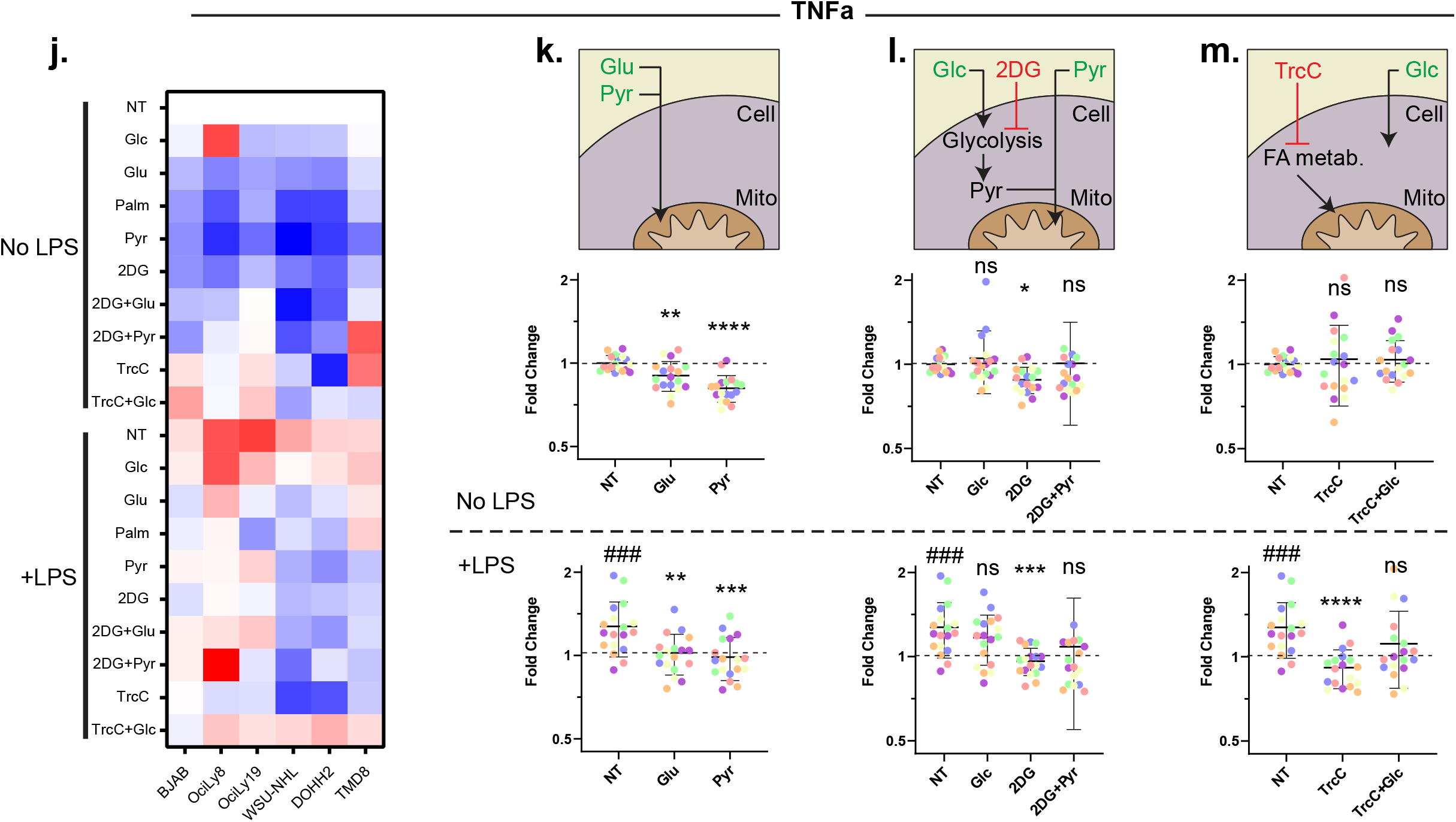
Effects of metabolic stimulation and inhibition on IL-6, IL-10 and TNFα production by DLBCL cell lines in vitro. Human DLBCL cell lines (BJAB, OciLy8, OciLy19, WSU-NHL, DOHH2, TMD8) were treated for 24 hours in full growth medium with metabolic supplements: glucose (Glc, 2mM), glutamine (Gln, 2mM) palmitate (Palm, 5µM), pyruvate (Pyr, 2mM) or metabolic inhibitors: 2-deoxy-d-glucose (2-DG, 5mM) and Triacsin C (TrcC, 0.1µM) with or without lipopolysaccharide (LPS, 5µg/mL) stimulation (**A)**. IL-6 (**B)**, IL-10 (F) and TNFα (J) production after 24-hour treatment was determined via ELISA and fold changes were plotted in a heatmap. Effects of mitochondrial substrates (**C, G, K)**, glycolytic modulation (**D, H, L)** or fatty acid inhibition (**E, I, M)** on IL-6, IL-10 and TNFα, respectively, were plotted in dot plots. Each dot represents an individual repeated experiment. Each colour represents the cell line. Data of three individual experiments is plotted. Two-way ANOVA with Tukey’s Post-Hoc test was performed to check for significant differences between the indicated groups, ns: not significant, *p<0.05, **p<0.01, ***p<0.001 ****p<0.0001, #### p<0.0001 compared to non-treated, no LPS. Experiments were repeated three times. Error bars represent mean +/-SD.

### Glycolysis pathway is important for in vitro IL-6 production by DLBCL cells

When supplemented with the different metabolic substrates and inhibitors, IL-6 production changed in a substrate-dependent manner (Figure 2B). On average, the cell lines significantly decreased their IL-6 production upon pyruvate supplementation but not when supplemented with glutamine, nor were these effects significant in LPS-treated cells (Figure 2C). Without LPS, glucose supplementation significantly increased IL-6 production, whereas inhibition of glycolysis by 2-DG significantly decreased production (Figure 2D). Pyruvate supplementation in combination with 2-DG was not sufficient to rescue the glucose-induced IL-6 production, indicating that glycolysis intermediates are important for IL-6 production, and not the end-product pyruvate. Inhibition of fatty acid metabolism by TriacsinC significantly reduced IL-6 production, which could be rescued by additional glucose supplementation (Figure 2E). In general, LPS stimulation did not affect the IL-6 production. However, it did abolish the glucose-induced IL-6 production and diminished metabolism-induced changes to IL-6 production in general. Nevertheless, even in combination with LPS, glycolysis inhibition by 2-DG decreased IL-6 production significantly, indicating that the glycolysis intermediates are important for in vitro IL-6 production.

### Glycolysis and fatty acid metabolism are important for in vitro IL-10 production in unstimulated DLBCL cells

Similar to the observed substrate-dependent IL-6 production, IL-10 production differed depending on metabolic supplementation or inhibition (Figure 2F). Single supplementation of mitochondrial substrates glutamine and pyruvate did not alter IL-10 production, independent of LPS stimulation (Figure 2G). In contrast, glucose supplementation in unstimulated DLBCL cells significantly increased IL-10 production (Figure 2H). Moreover, inhibition of glycolysis drastically decreased IL-10 production. Pyruvate supplementation in combination with glycolysis inhibition could not rescue IL-10 production, suggesting the potential involvement of glycolysis intermediates. Finally, in unstimulated DLBCL cells, inhibition of fatty acid metabolism by TriacsinC significantly decreased IL-10 production (Figure 2I). Strikingly, this could not be rescued by addition of glucose to TriacsinC, potentiating the idea that both fatty acid metabolism and glycolysis are important to sustain normal IL-10 production in vitro. When adding LPS, these patterns were no longer as clearly displayed. Some cell lines increased overall IL-10 production upon LPS stimulation (mainly OciLy8 and OciLy19), even overcoming 2-DG-and TriacsinC-induced decrease in IL-10 production, suggesting that in these cells, the LPS-TLR4 cascade is dominant over metabolic-induced changes. In general, the same trend for glycolysis-dependent changes in IL-10 production was still present in LPS-stimulated DLBCL cells. However, due to the huge spread between cell lines, it is clear that TLR4 activation does alter metabolic preferences and subsequent IL-10 production differently in these cell lines. In summary, these results suggest a role for glycolysis and fatty acid metabolism to maintain IL-10 production, independent on stimulation.

### Metabolic balance is essential for in vitro TNFα production in DLBCL cells independent of LPS

In line with the IL-6 and IL-10 data, metabolic-treatment-dependent effects on the in vitro TNFα production in DLBCL cells were observed (Figure 2J). Interestingly, supplementation of mitochondrial substrates glutamine and pyruvate, both significantly decreased TNFα production (Figure 2K), suggesting that mitochondrial activation or TCA-intermediates hamper with the TNFα cascade. Glucose supplementation did not affect TNFα production (Figure 2L). This suggests that glucose was not completely broken down into pyruvate because that should have resulted in a decrease similar to pyruvate supplementation, which can be readily taken up by the cells. Simultaneously, the absence of glucose-mediated increase in TNFα also suggests that glycolysis intermediates are not essential for TNFα production. Although inhibition of glycolysis did decrease TNFα production, pyruvate in combination with 2-DG could rescue normal TNFα production, suggesting the concentration of pyruvate is essential, with too much or too few molecules pyruvate both leading to reduced TNFα. Future research should involve concentration gradients of pyruvate to test this hypothesis. Moreover, LPS-stimulation alone significantly increased TNFα production in DLBCL cells, without any other metabolic supplementation (Figure 2K). Furthermore, LPS in combination with mitochondrial substrates significantly abolished the LPS-induced increase in TNFα production. Similarly, glucose in combination with LPS did not affect the production beyond the LPS-induced effect, whereas 2-DG did abolish the TNFα production and combination with pyruvate could again rescue the production. Finally, even though FA-metabolism inhibition by TriacsinC alone did not affect TNFα, it did abolish the LPS-induced TNFa production, yet this could be rescued by additional glucose supplementation (Figure 2M). Since both glycolysis inhibition and FA-inhibition could be rescued by substrate supplementation, these results suggest a positive LPS-dependent regulation of TNFα that is likely partly dependent on mitochondrial metabolic balance.

### In vitro metabolism-dependent IL-6 and IL-10 production is similar between all unstimulated DLBCL cells, TNFα most similar between stimulated DLBCL cells

In the previous section we showed that metabolic supplementation or inhibition resulted in different effects on the IL-6, IL-10 or TNFα production by the DLBCL cell lines. Unfortunately, by pooling the different cell lines, subtle differences between cell lines are lost. To illustrate, we observed an overall significant increase in IL-10 production after glucose supplementation, and an overall decrease in IL-10 production after 2-DG inhibition (Figure 2H). However, this does not show whether the different cell lines displayed comparable increase/decrease patterns based on treatment. This is important for the interpretation of the data, since a ‘universal’ shared response pattern would indicate more conserved, and therefore physiologically more relevant, mechanisms of interaction between metabolism and cytokine production. Therefore, we next assessed if the metabolism-dependent cytokine profiles were cell line specific or broadly applicable to DLBCL cell lines. To this end, the response of each cell line to every treatment was plotted against the response of every other cell (Figure 3B). Through the datapoints, the correlation coefficient (*R*-value) could be determined with an associated chance (p-value). The *R*-value indicates whether the two cell lines plotted against each other had a similar response pattern over the different treatments (positive *R*-value), and the p-value indicates how well the datapoints of each treatment stick to the fitted line (Figure 3B). Furthermore, by correlating the effect per treatment on one cytokine (i.e., IL-6), with the effect per treatment on another cytokine (i.e., IL-10), potential shared metabolic pathways between cytokines could be identified (Figure 3C). By combining the *R*-values in a correlation matrix, each cell-cell correlation could be visualized for each cytokine. For example, unstimulated BJAB and OciLy8 had similar IL-6 production in response to the different treatments and correlated positive, *r*(8) = .77, p=0.0095, where *r* is Pearson’s R, (8) is the degrees of freedom, .77 is the *r*-value and p=0.0095 the chance that this correlation occurred by chance, with everything below p=0.05 deemed significant (Figure 3D). Furthermore, the IL-6 production and TNFα production of unstimulated BJABs was observed not similar, with a negative correlation, *r*(8) = -.15, p=.67. In the correlation matrix, each comparison is indicated by a colour corresponding to its *r*-value (Red=max: 1; white=median: 0.19, blue=min: - 0.61). The exact *R*-values and associated p-values per comparison can be found in the supplementary material.

**Figure 3.**
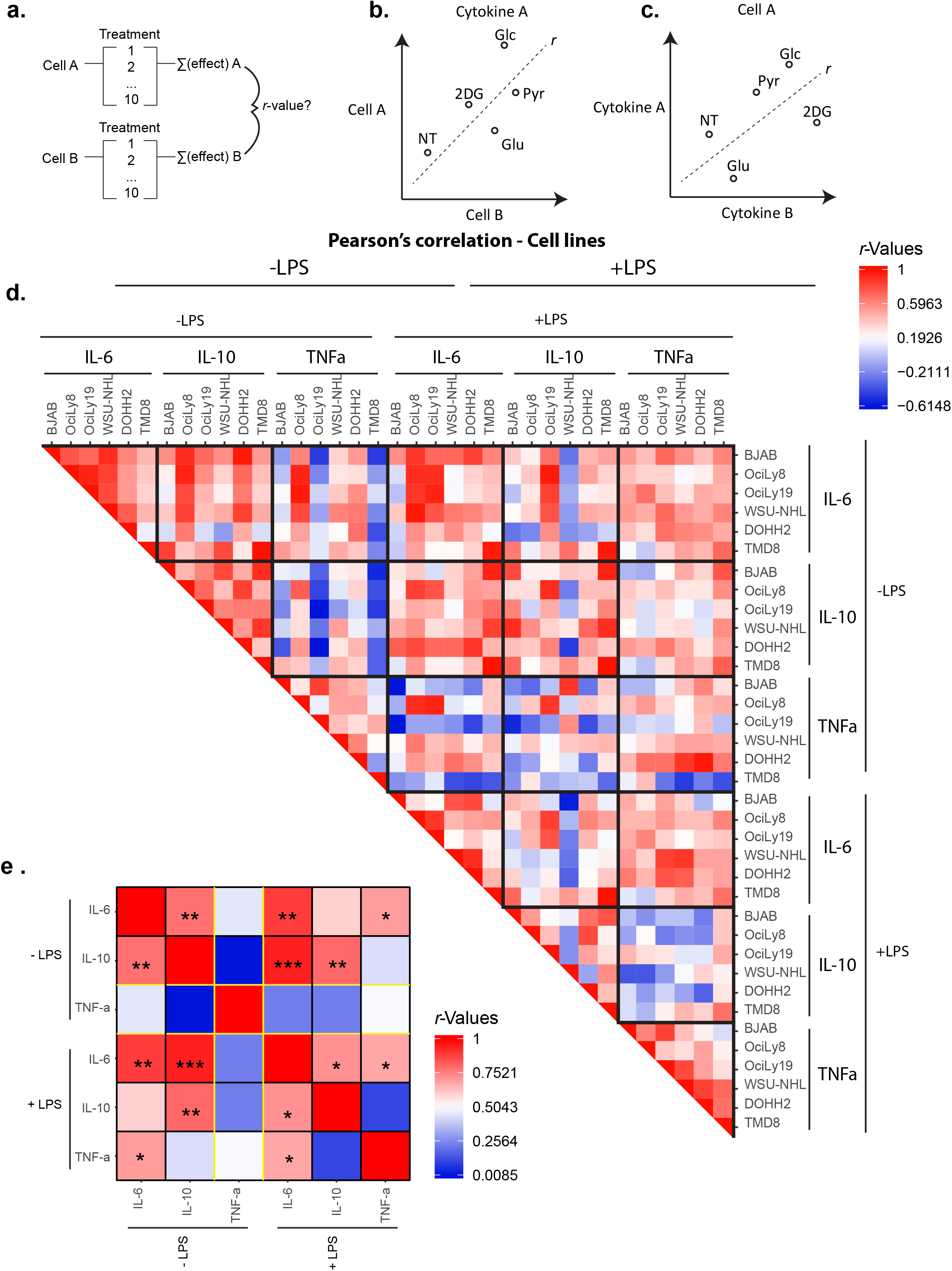
Pearson’s correlation revealed shared metabolism-dependent cytokine-response patterns between cell lines. Data from figure 2 was used to determine correlation between cell lines per treatment. For each cell-cytokine combination, the effect of each treatment on the cytokine production was correlated to the effect of each treatment on the cytokine production of any other cell (**A)**. Cartoon depicting how the data was used to determine correlation between cell lines (*R*-value) (**B)**. Each treatment had an effect on the cytokine production of cell A and cell B. These effects were plotted as x and y coordinates for cell A and B, respectively. After which a best fit line would determine the correlation (positive *r*: positive correlation, cell line A and B responded similar to the metabolic treatments; low *r*: no correlation, cell line A and B did not respond similar to the metabolic treatments). Cartoon depicting how the data was used to determine correlation between cytokines (**C)**. Each treatment had an effect of cytokine A and cytokine B on cell A. These effects were plotted as x and y coordinates for cytokine A and B, respectively. A best-fit line determined the correlation. The spread in datapoints from the best-fit line determined the chance that the correlation was based on chance (p-values smaller than 0.05 were deemed significant, not based on chance) (**C)**. *R*-values for each cell-treatment-cytokine combination is displayed in a heatmap (Red=max: 1; white=median: 0.02, blue=min: -0.97) (**D)**. *R*-values for the average (of the six cell lines) treatment effect on each cytokine IL-6, IL-10 and TNFα is displayed in a heatmap (Red=max: 1, white=median: 0.504, blue=min: 0.0085) (**E)**. Experiments were repeated three times. The exact *R*-values and associated p-values per comparison can be found in the supplementary material. *p<0.05,**p<0.01, ***p<0.001.

Without LPS stimulation, we observed a strong positive correlation between the cell lines in their IL-6 response, as well as in their IL-10 response, with an average *r*(8)= .67 and *r*(8) = .68 respectively (Figure 3D). In contrast, TNFα production correlated only moderately positive between cell lines, average *r*(8) = .49. LPS stimulation reduced the correlation between cell lines in their IL-6, and IL-10 production, with an average *r*(8)= .55 and *r*(8) = .45 respectively. In contrast, LPS stimulation drove the different cell lines to a comparable TNFα production, illustrated with a positive correlation, average *r*(8) = .57. These results suggest that fluctuations in IL-6 production and IL-10 production in response to metabolic manipulations are not cell-line specific and might apply to DLBCL in a general sense, whereas LPS mostly attenuates TNFα production in response to metabolic fluctuations.

After confirmation of positive correlation between cell lines in their response to the different metabolic treatments, the overall similarities or differences between cytokine responses were analysed. To this end, the average cytokine-response over all the cell lines, to each treatment, was plotted in a correlation matrix (Figure 3E). This allowed us to see whether the different cytokines, were produced similarly in response to the metabolic treatments. Without LPS stimulation, IL-6 and IL-10 correlated positively, *r*(8) = .78, p = .0079. In contrast, TNFα correlated only minimally positive to IL-6, *r*(8) = .46, p = .18 and did not correlate to IL-10 at all *r*(8) = .01, p = .98. In contrast, in LPS-stimulated DLBCL cells, the in vitro IL-6 production correlated positively to TNFα production, *r*(8) = .68, p=0.0299. The correlation between IL-6 and IL-10 in LPS stimulated DLBCL cells was similar to the unstimulated situation, *r*(8) = .72, p=0.0200. Similarly, TNFα and IL-10 also did not correlate in LPS stimulated DLBCL cells, *r*(8) = .11, p=0.75. These results suggest that the IL-6 and IL-10 production of unstimulated DLBCL cells respond similar to metabolic manipulations and might have shared pathways. Furthermore, it suggests that LPS stimulation drives DLBCL cells to produce TNFα and IL-6 in response to similar metabolic manipulations, but not TNFα and IL-10.

### Mitochondrial substrate supplementation and glycolysis inhibition define cytokine profile of DLBCL cells

We next assessed how the different metabolic manipulations correlated to each other and how much they caused similar or dissimilar responses between cytokines. Here, the effect of each treatment on the different cell lines was correlated to the effect of the other treatments on the cell lines (Figures 4A, B). This visualized whether different treatments had similar effects and whether same treatments had similar effects on different cytokines. This means that if two treatments resulted in similar cytokine production, they would correlate positively. Similarly, if one treatment had a comparable effect on two different cytokines, they would correlate positively, indicated by an *R*-value close to 1 and a red colour in the correlation matrix (Red=max: 1; white=median: 0.02, blue=min: -0.97). The exact *R*-values and associated p-values per comparison can be found in the supplementary material.

**Figure 4.**
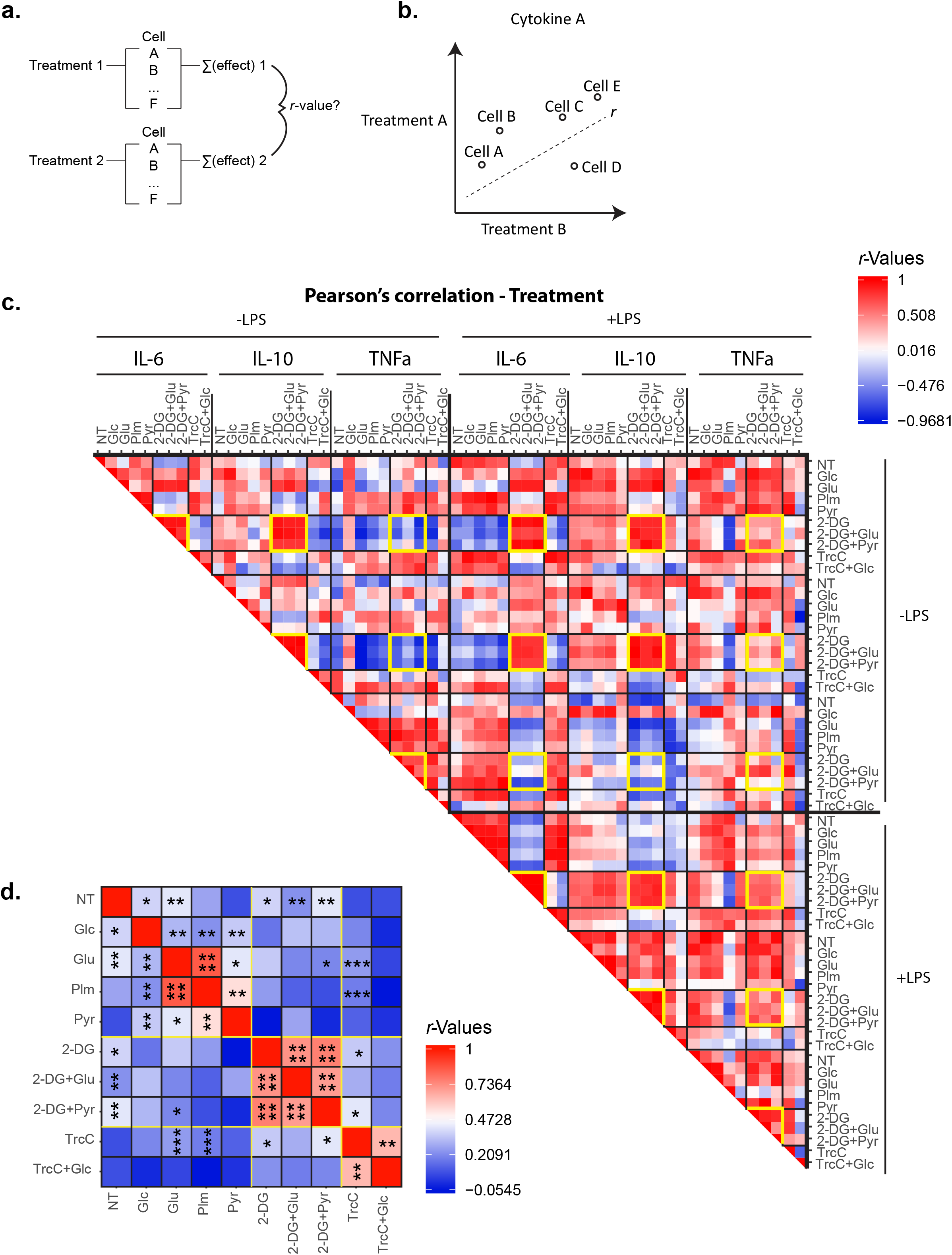
Pearson’s correlation revealed shared and dissimilar cytokine-response patterns between metabolic treatments. Data from figure 2 was used to determine correlation between treatments per cytokine. For each treatment-cytokine combination, the effect on each cell line was plotted against the effect of another treatment on the same cells (**A)**. Cartoon depicting how the data was used to determine correlation between cell lines (*R*-value) (**B)**. Treatment 1 had a certain cytokine-effect on cell A-F, and treatment 2 also had a certain cytokine-effect on cell A-F. These effects were plotted as x and y coordinates for treatment A and B, respectively. A best fit line would determine the correlation (positive *r*: positive correlation, cells A-F responded similar to the metabolic treatments 1 and 2; low *r*: no correlation, cell lines A-F did not respond similar to the metabolic treatments 1 and 2). The spread in datapoints from the best-fit line determined the chance that the correlation was based on chance (p-values smaller than 0.05 were deemed significant, not based on chance) (**B)**. *R*-values for each treatment-cytokine combination is displayed in a heatmap (Red=max: 1; white=median: 0.02, blue=min: -0.97, yellow squares indicate correlation between glycolysis inhibition treatments) (**C)**. *R*-values for the average (of the six cytokines) treatment effects on the combined cytokine production is displayed in a heatmap (Red=max: 1, white=median: 0.47, blue=min: -0.05) (**D)**. Experiments were repeated three times. The exact *R*-values and associated p-values per comparison can be found in the supplementary material. See also Figure S1. *p<0.05, **p<0.01, ***p<0.001, * * * * <p<0.0001.

A striking positive correlation was observed between glutamine, palmitate and pyruvate supplementation in how they affected IL-6 and TNFα production in unstimulated DLBCL cells, with an average r(4) = .67 and r(4) = .88, respectively (Figures 4C S1). In LPS-stimulated DLBCL cells, these positive correlations among the mitochondrial substrates were even stronger for IL-6 (r(4) = .94), and also for IL-10 (r(4) = .82). In contrast, LPS stimulation abolished the correlation among the mitochondrial substrates in their effect on TNFα production (r(4) = .42). Furthermore, we observed a striking correlation in glycolysis inhibited DLBCL cells, both in unstimulated and stimulated situations (Figure 4C, indicated with the yellow lines). In unstimulated DLBCL cells, 2-DG strongly correlated with 2-DG in combination with glutamine or pyruvate. This suggests that most of the effect caused by glycolysis inhibition was indeed caused by absence of glycolysis intermediates, and not by lack of pyruvate, since that would cause a rescue and therefore a negative correlation to 2-DG alone. Interestingly, the effect of 2-DG (with or without additional pyruvate or glutamine) was very similar on IL-6 production and IL-10 production (r(4) = .91, p=0.0107) but very dissimilar to the effect it had on TNFα production (r(4) = -.87, p=0.0261). However, 2-DG-induced changes to TNFα were more similar to IL-6 changes in LPS-activated cells (r(4) = .56, p=0.24). These results suggest a dependency on glycolysis of DLBCL cells for their IL-6 and IL-10 production in unstimulated situations, whereas this only applies to TNFα in LPS-activated DLBCL cells.

We next analysed the effect sizes of the different metabolic treatments on all the other variables. This would provide insight into which treatments were most determining factors in the cytokine profiles of DLBCL. Importantly, over all the different cytokines and cell lines, non-treated cells (NT) did not correlate with any other treatment, confirming what we observed before, that metabolic treatments really altered cytokine profiles (Figure 4D). Glucose supplementation correlated with no other treatment, suggesting that its effect on cytokine production is unique and likely due to glycolytic derivates instead of mitochondrial derivates, as this would result in a positive correlation with pyruvate. Furthermore, over all variables, mitochondrial supplements glutamine and palmitate correlated strongest, r(34) = .88, p<0.0001. Glycolytic inhibition proved very dominant in determining cytokine profiles, as it strongly correlated with glycolytic inhibition in combination with glutamine and pyruvate, but not with any other treatment (2-DG+glu; r(34) = .88, p<0.0001, 2-DG+pyr; r(34) = .81, p<0.0001). Finally, FA-metabolism was observed to be of importance, since inhibition of FA did correlate with FA-inhibition in combination with glucose, but not with any other treatment, r(34) = .47, p=0.0035, suggesting the difference in cytokine production could not be rescued. In summary, these results confirm earlier observations and identify mitochondrial supplementation, glycolytic inhibition and FA-inhibition as important factors in the cytokine production of DLBCL.

### Exploring metabolic response to IL-4 in germinal centre B- and activated B-cell-DLBCL subsets

The previous results suggest that cytokine release by lymphoma cells can be modulated by metabolic pathways like glycolysis. If these cytokines, in turn, signal other cells to alter their metabolism, this could theoretically lead to a propagating domino effect. It seems unlikely that any physiological situation will be so easily destabilized. Nevertheless, investigating whether metabolism is involved in positive feedback loops in cytokine pathways may help us to better understand how cytokine-associated pathologies occur. Illustrative of this was the finding of metabolism-cytokine crosstalk, that drove the cytokine profile of COVID-patients, and it was explored as a viable therapeutic intervention strategy against COVID-induced cytokine storms^13,38^. In fact, aberrant glucose metabolism explained why diabetic patients with COVID were overrepresented in the ICUs^39^. Therefore, we explored whether the metabolic phenotype of a lymphoma cell changes upon cytokine stimulation. Established cell-models were selected that represent two subtypes of DLBCL, germinal centre B-cell (GCB-) DLBCL and activated B-cell (ABC-) DLBCL. We chose the two subsets in an attempt to distinguish whether any potential metabolic shift caused by cytokines, could be disease-specific, or represent a more universal cytokine-mediated response. Furthermore, autocrine signalling had to be excluded in order to study whether any metabolic shift really was caused by a newly introduced cytokine. Therefore, production of several cytokines was measured by two GCB-DLBCL lines (WSU-NHL and OciLy8) and three ABC-DLBCL (HBL1, U2932 and TMD8). IL-4 was not produced by any of the cell lines (Figure S2). Interestingly, it is known that GCB-DLBCL and ABC-DLBCL have a distinct response to IL-4, with the former becoming highly proliferative and increasingly sensitive to immuno-chemotherapy upon IL-4 supplementation^40^.

### GCB-DLBCL increased mitochondrial metabolism and ABC-DLBCL increased glycolysis metabolism after in vitro IL-4 stimulation

The metabolic phenotypes in response to IL-4 were assessed by continuous measurement of oxygen consumption rate (OCR) and extracellular acidification rate (ECAR) in response to mitochondrial and glycolytic inhibitors (Figures 5A, J). OCR as proxy for mitochondrial activity, and ECAR as proxy for glycolytic activity, provide insights into metabolic activity of the different DLBCL-subsets. Sequential administrating of mitochondrial inhibitors and uncouplers to two GCB-DLBCL cell lines (Figures 5B, C) and three ABC-DLBCL cell lines (Figures 5D, E and F), revealed that GCB-DLBCLs increased their basal respiration (Figure 5G), ATP production (Figure 5H) and spare respiratory capacity (SRC) (Figure 5I) significantly upon IL-4 stimulation, whereas no IL-4-induced mitochondrial response was observed in the ABC-DLBCLs. After submitting the DLBCL subtypes (Figures 5K-O) to glycolytic stimulation and inhibition, the effect of IL-4 on the basal glycolytic activity and the glycolytic reserve was determined. IL-4 did not affect the basal glycolytic activity of either DLBCL subtype (Figure 5P). In contrast, the glycolytic reserve of the ABC-DLBCL subtype increased significantly upon IL-4 stimulation, whereas the glycolytic reserve of GCB-DLBCL cells remained unaffected (Figure 5Q). These results suggest that specifically GCB-DLBCLs increase their mitochondrial breakdown of substrate when stimulated with IL-4. The ATP-associated oxygen consumption suggests, that GCB-DLBCLs used the enhanced mitochondrial activity to synthesize ATP and provide energy. Even though the GCB-DLBCL increased their mitochondrial respiration upon IL-4 stimulation, they still displayed an SRC, indicating they possessed surplus TCA intermediates. Although the SRC of ABC-DLBCL was not significantly increased upon IL-4 stimulation, cell-line specific differences were found as two out of three (HBL1 and U2932) lines did show increased SRC. Furthermore, ABC-DLBCL seemed more glycolytically active after IL-4 stimulation. In conclusion, the results suggest a disease-specific metabolic response to exogenous cytokines.

**Figure 5.**
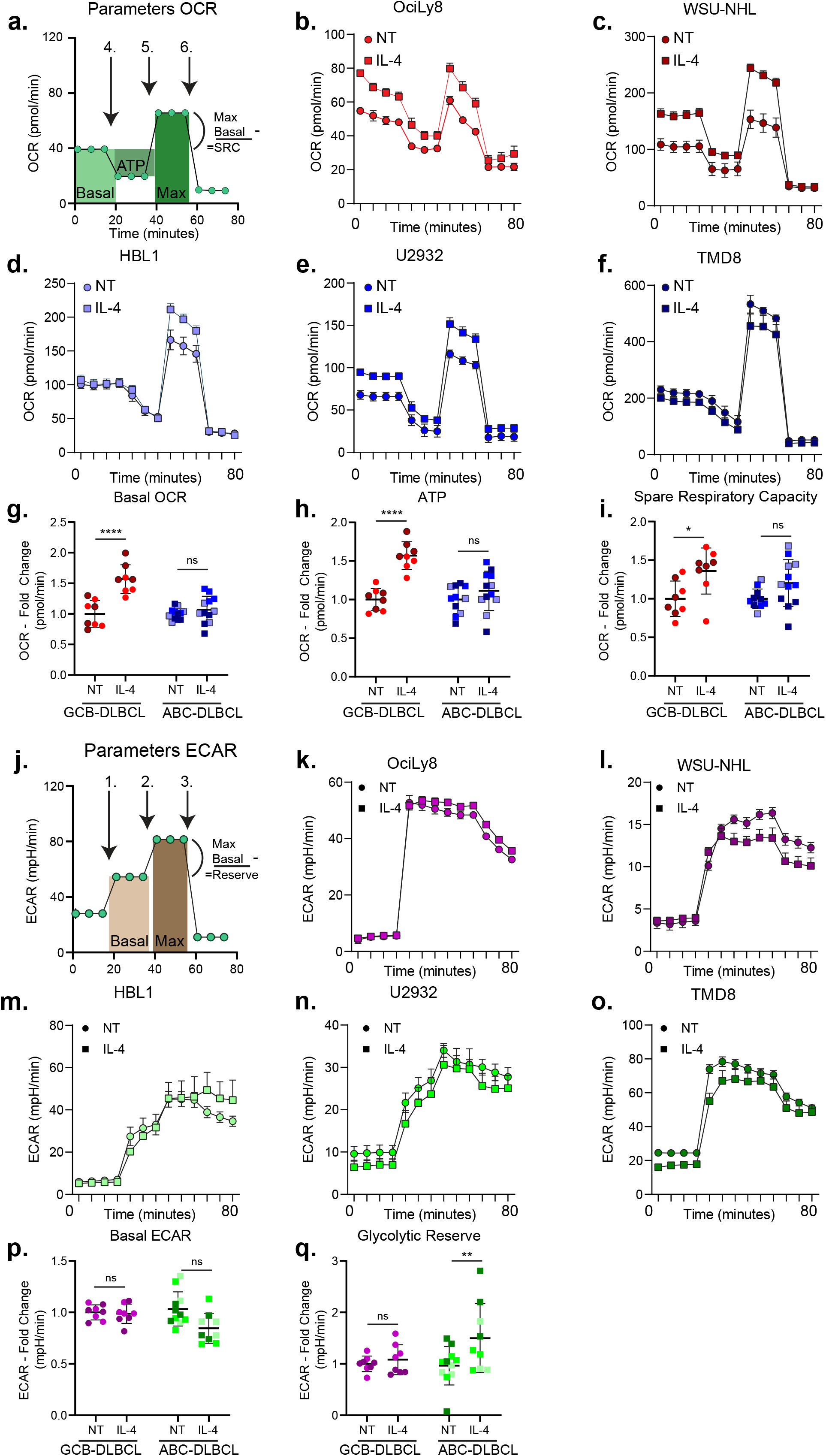
Different metabolic preferences of germinal centre-B and activated-B cell-diffuse large B-cell lymphoma cell lines in vitro. Seahorse analysis of mitochondrial (**A)** oxygen consumption rate (OCR) in response to ATP-synthase inhibition with oligomycin A (**1**, 1µM), mitochondrial uncoupling with FCCP (**2**, 1µM) and complex I and II inhibition with rotenone/antimycin A (**3**, 1 µM). Continues OCR measurements of germinal centre-B (GCB) diffuse large B-cell lymphoma (DLBCL) cell lines (**B, C)** and activated B-cell DLBCL (**D-F)** in response to 2 hours IL-4 stimulation. Basal OCR (**G)**, ATP (**H)** and spare respiratory capacity (SRC) (**I)** were calculated as indicated in **A a**nd displayed in fold change relative to non-treated cells (NT). Continuous glycolytic extracellular acidification rate (ECAR) (**J)** in response to glucose injection (**4**, 10mM) mitochondrial inhibition with oligomycin A (**5**, 1µM), glycolysis inhibition with 2-deoxyglucose (2-DG) (**6**, 20 mM). Basal ECAR (**P)** and glycolytic reserve (**Q)** were calculated as indicated in J and displayed in fold change relative to NT. See also Figure S2. Two-way student T-tests between NT and IL-4 stimulated was performed to check for significant differences between the indicated groups. *p<0.05,**p<0.01, * **p<0.001, * * * * <p<0.0001.

## Discussion

Aberrant cytokine production by B cell lymphomas has been associated with inferior prognosis and inferior therapeutical response. The notion that metabolism affects tumour cell function has become commonly accepted over the past few years. However, it remains unclear if and how metabolic pathways control cytokine production in (B-cell) tumours. Here, we show the potential of metabolic manipulations to attenuate IL-6, IL-10 and TNFα production by DLBCL cells. Furthermore, we show that exogenous IL-4 affects the metabolic phenotype of two DLBCL subtypes differentially.

Glycolysis activating kinases^41^ or glycolytic enzyme quantities^42^ are both prognostic markers for significantly worse disease progression in DLBCL. In fact, inhibition of glycolysis might prove a valid therapeutical strategy against DLBCL^30,42^. Furthermore, it is well established that elevated serum levels of cytokines, especially IL-6 and IL-10, are observed in DLBCL patients and correlate with worse prognosis^43,44^. New insights in the role of metabolism in cytokine production, could lead to better therapeutical options for DLBCL since both IL-6 and TNFα are important cytokines in DLBCL. IL-6 is a B cell growth factor^45^ and essential for GC formation^37^ and response^46^. Furthermore, it is heavily associated with lymphomagenesis in DLBCL^31,47^, correlated to worse overall survival (OS), and linked to therapy resistance^32^. IL-6 and TNFα are both under control of the JAK2/STAT3 pathway^48^ and are both associated with worse prognosis in DLBCL^49^, as well as in autoimmunity^50^. Furthermore, IL-6 works synergistically with TNFα^51–54^, which on its own is associated with B cell development, lymphomagenesis, and therapy resistance and tumour-promoting actions in carcinogenesis^55^. Our results suggest that glycolysis-targeting strategies in DLBCL can be worth investigating further. Potential therapies could work two-fold, both by limiting energy and building block maintenance in the highly proliferative lymphoma cells, as well as limiting cytokine production via glycolytic intermediates.

The mechanisms underlying glycolytic intermediate involvement in cytokine production remain unknown for now. Recently, lactate, the end-point of glycolysis, was shown to modify histones of LPS-activated macrophages, thereby allowing for targeted gene expression associated with an inflammatory phenotype^56,57^. Targeted histone modifications may affect cytokine-associated gene-expression in glycolytic DLBCL. Alternatively, aberrant activity of glycolytic enzymes could explain part of the cytokine response. GAPDH for example, was shown to bind to and suppress translation of certain mRNAs, including that of cytokines^58,59^. Since 2-DG blocks the first step of the glycolysis pathway, GAPDH, which is more downstream in the cascade would be less engaged by glycolytic intermediates and as such could bind more mRNAs, thereby blocking translation. In contrast, addition of glucose would lead to more substrate to engage GAPDH and as such would limit its translation-blocking capacity. This could explain our results, although it seems likely these mechanisms are at least partially cell-type specific, as intraperitoneally injection of 2-DG in mice was found to increase both TNFα and IL-6 concentrations in blood^11^. It would therefore be beneficial to study these principles in healthy B cells and in primary B cell malignancies. Inhibition studies combined with quantitative PCR and cytokine-vectors either with or without the AU-rich elements could shine light on the matter.^60,61^ If GAPDH indeed controls IL-6, IL-10 and TNFα production by binding to AU-rich elements of cytokine-mRNA, the vectors without the AU-rich elements should always be actively engaged and translated, whereas the AU-rich element-containing vectors would fail to do so in a 2-DG dependent manner. If neither would be affected, it is more likely that histone lactylation or other metabolic influences would control the cytokine production^62^. In that case, Chromatin immunoprecipitation (ChIP) would provide insight into the involvement of the promotor regions of the cytokine-associated genes during glycolysis inhibition.

Our results also suggest an involvement of mitochondrial substrates in limiting IL-6 and TNFα production. Recent studies have convincingly shown a role of TCA intermediates in immune-modulation^63–65^. Among others, citrate and fumarate modulate histone acetylation and demethylation, respectively, whereas α-ketoglutarate inhibits hypoxia-induced factor 1α (HIF1α), a well-known glycolytic modulator. Nevertheless, the total picture is still far from complete, with many intermediates exerting multiple functions, sometimes dependent on the cell-type, sometimes dependent on the cell-fate and sometimes dependent on the concentrations of transporters and other TCA-intermediates^66^. Moreover, the contribution of secondary mitochondrial products such as mitochondrial reactive oxygen species (mtROS) and mitochondrial DNA (mtDNA) further complicate the picture^67,68^. Follow-up studies with more aimed metabolites could help clarify the story in DLBCL. In the present study, we provided the cells with substrate that could be used to fully cycle through the TCA-cycle and could lead to a build-up and subsequent effect of either one of the TCA-intermediates. Follow-up studies should involve a more elaborate supplementation profile, where the cells are supplemented with specific TCA intermediates that potentially regulate immune cell functioning. Combining these studies with inhibitors for mitochondrial-substrate carriers would provide insight into whether the added substrates exert a cytosolic function. Mitochondrial citrate for example, could be used as substrate in the TCA-cycle, thereby generating alpha-ketoglutarate (a-KG) which in turn is involved in histone modifications^69^. In contrast, cytosolic citrate is involved in *de novo* lipid synthesis and is an activator of ATP-citrate lyase (ACLY), both of which can in turn attenuate the inflammatory profile of an immune cell^70–72^. By selectively providing the different substrates one by one and measuring cytokine production, we could pinpoint where the effects are caused.

Lastly, our results suggest a different metabolic response to IL-4 stimulation in GCB- and ABC-DLBCL, with GCB-DLBCL displaying increased mitochondrial activity and ABC-DLBCL becoming more glycolytically flexible. IL-4 has been reported to have different effects on different DLBCL subtypes^73,74^ but underlying mechanism have not been fully resolved. Our data suggests that GCB-DLBCL become metabolically much more activated and could potentially generate more energy, more TCA-intermediates, and more building blocks to facilitate rapid proliferation when stimulated with IL-4. This could explain why IL-4 sensitizes GCB-DLBCL to doxorubicin (Dox), as Dox is known to permeabilize mitochondria, therefore being more toxic for mitochondrial-dependent proliferative cells^75^. Further research is required to determine if ABC-DLBCLs would be better treated with glycolytic inhibitors in combination with IL-4. In conclusion, our work shows that there is a connection between cellular metabolism and the cytokine response in DLBCL in vitro. It paves the road for more in-depth studies to potentially reveal metabolic targets for therapeutic intervention where both proliferative capacity and survival-stimulating cytokines can be abolished.

## Supporting information

Supplemental text figure 1 ang 2

## Acknowledgements

R.P. and A.v.S. conceived the idea for this project. R.P. conducted the experiments and wrote the article with valuable input of A.v.S. and S.v.D. This work is supported by the European Research Council (ERC Consolidator Grant 724281) and Dutch Research Council (NWO, OCENW.XS25.3.426).

## Materials and Methods

### Human cell lines

Human lymphoma BJAB (Cat: ACC 757), OciLy8, OciLy19 (Cat: ACC 528), WSU-NHL (Cat: ACC 58), DOHH2 (Cat: ACC 47), TMD8 (Cat: CVCL_A442), U2932 (Cat: ACC 633) and HBL1 (Cat: CVCL_4213) cells were cultured in RPMI1640 medium supplemented with 10% heat-inactivated foetal bovine serum (FBS), 1% antibiotic-antimycotic (AA) and 1% ultra-glutamine (UG) (Gibco, Thermo Fisher Scientific, Waltham, MA, USA) and maintained at 37°C with 5% CO_2_ . Cells were seeded at ∼3e^5^ cells/mL at the day prior to experiments. Cell lines were routinely checked for absence of mycoplasma contaminations. Cell lines were derived from DSMZ, ATCC and Dr. Blanca Scheijen (Dept. Pathology, Radboudumc) and cell lines were authenticated using STR-analysis.

### Cytokine quantification studies

Cells were seeded at ∼3e^5^ cells/mL in 200µL culture medium plus the indicated metabolic substrates or inhibitors in 96-well flat-bottom plates (Corning Incorporated, New York, NY, USA) and kept at 37°C with 5% CO_2_ overnight. Substrates glucose, glutamine, pyruvate (Gibco, Thermo Fisher Scientific, Waltham, MA, USA), palmitate (Agilent Technologies, Santa Clara, CA, USA) and inhibitors 2-deoxyglucose (2-DG), Triacsin C (Abcam, Cambridge, UK) were diluted and stored in accordance with manufacturers advice. After 24 hours of treatment, cells and supernatants were transferred to 96-well V-bottom plates (Corning Incorporated, New York, NY, USA) and spun down at 1500 RPM for 3 minutes. Supernatants were collected and transferred to primed ELISA plates according to manufactures protocols (IL-4, Cat: # KHC0041; IL-6, Cat: # EH2IL6; IL-10, Cat: # KHC0101; TNFα, Cat: # KHC3011). Signal strength was determined with Bio-Rad iMark microplate absorbance reader (Bio-Rad, Hercules, CA, USA).

### Seahorse metabolic analyser studies

Metabolic respiratory assays were performed as described by the Seahorse manufacturer (Agilent Technologies, Santa Clara, CA, USA). Cells were treated Cells were seeded on Cell-Tak (Corning Incorporated, New York, NY, USA) coated Seahorse cell culture plates and exposed to Seahorse XF Base Medium supplemented with 2 mM glutamine (For measuring extracellular acidification rate, ECAR), or 1 mM pyruvate, 2 mM glutamine and 10 mM glucose (For measuring oxygen consumption rate, OCR). Cells were subsequently rested in 0% CO_2_ at 37°C for 45-60 minutes before measuring.

### Statistical analysis

Statistical analysis was performed with GraphPad Prism 8. Performed tests, number of individual replicates and sample sizes are described in figure legends. Exact p-values can be found in supplementary data.

## References

1. Dinarello, C. A. Historical Review of Cytokines. Eur. J. Immunol. 37, S34 (2007).

2. Leonard, W. J. Cytokines and immunodeficiency diseases. Nat. Rev. Immunol. 1, 200–208 (2001).

3. Moudgil, K. D. & Choubey, D. Cytokines in Autoimmunity: Role in Induction, Regulation, and Treatment. J. Interf. Cytokine Res. 31, 695 (2011).

4. Gupta, M. et al. Elevated serum IL-10 levels in diffuse large B-cell lymphoma: a mechanism of aberrant JAK2 activation. Blood 119, 2844–2853 (2012).

5. Stenson, P. D. et al. Human Gene Mutation Database (HGMD): 2003 update. Hum. Mutat. 21, 577–81 (2003).

6. Tabatabaei-Panah, P. S. et al. Proinflammatory cytokine gene polymorphisms in bullous pemphigoid. Front. Immunol. 10, 636 (2019).

7. Tarique, M. et al. Association of IL-10 Gene Polymorphism With IL-10 Secretion by CD4 and T Regulatory Cells in Human Leprosy. Front. Immunol. 11, 1974 (2020).

8. Medzhitov, R. & Horng, T. Transcriptional control of the inflammatory response. Nat. Rev. Immunol. 9, 692–703 (2009).

9. ater Horst, R. et al. Host and environmental factors influencing individual human cytokine responses Summary. Cell 167, 1111–1124 (2016).

10. Netea, M. G. et al. Defining trained immunity and its role in health and disease. Nat. Rev. Immunol. 2020 206 20, 375–388 (2020).

11. Dréau, D., Morton, D. S., Foster, M., Fowler, N. & Sonnenfeld, G. Effects of 2-deoxy-D-glucose administration on cytokine production in BDF1 mice. J. Interferon Cytokine Res. 20, 247–255 (2000).

12. Chiba, S. et al. Glycolysis regulates LPS-induced cytokine production in M2 polarized human macrophages. Immunol. Lett. 183, 17–23 (2017).

13. Xiao, N. et al. Integrated cytokine and metabolite analysis reveals immunometabolic reprogramming in COVID-19 patients with therapeutic implications. Nat. Commun. 2021 121 12, 1–13 (2021).

14. Wofford, J. A., Wieman, H. L., Jacobs, S. R., Zhao, Y. & Rathmell, J. C. IL-7 promotes Glut1 trafficking and glucose uptake via STAT5-mediated activation of Akt to support T cell survival IL-7 promotes Glut1 trafficking and glucose uptake via STAT5-mediated activation of Akt to support T cell survival. Blood 111, 2101–2112 (2007).

15. Versatile functions for IL-6 in metabolism and cancer. Trends Immunol. 36, 92–101 (2015).

16. Ganeshan, K. & Chawla, A. Metabolic Regulation of Immune Responses. Annu. Rev. Immunol. 32, 609 (2014).

17. Bodhale, N. et al. Cytokines and metabolic regulation: A framework of bidirectional influences affecting Leishmania infection. Cytokine 147, (2021).

18. D’Autréaux, B. & Toledano, M. B. ROS as signalling molecules: mechanisms that generate specificity in ROS homeostasis. Nat. Rev. Mol. Cell Biol. 8, 813–824 (2007).

19. Schmidl, C., Delacher, M., Huehn, J. & Feuerer, M. Epigenetic mechanisms regulating T-cell responses. J. Allergy Clin. Immunol. 142, 728–743 (2018).

20. Lau, C. M. et al. Epigenetic control of innate and adaptive immune memory. Nat. Immunol. 19, 963–972 (2018).

21. Rodriguez, A. E. et al. Serine Metabolism Supports Macrophage IL-1β Production. Cell Metab. 29, 1003–1011.e4 (2019).

22. Ariav, Y., Chng, J. H., Christofk, H. R., Ron-Harel, N. & Erez, A. Targeting nucleotide metabolism as the nexus of viral infections, cancer, and the immune response. Sci. Adv. 7, 6165 (2021).

23. Lin, H. M. et al. Relationship between Circulating Lipids and Cytokines in Metastatic Castration-Resistant Prostate Cancer. Cancers (Basel). 13, (2021).

24. Wang, B. et al. Metabolism pathways of arachidonic acids: mechanisms and potential therapeutic targets. Signal Transduct. Target. Ther. 2021 61 6, 1–30 (2021).

25. Millet, P., Vachharajani, V., McPhail, L., Yoza, B. & McCall, C. E. GAPDH Binding to TNF-α mRNA Contributes to Posttranscriptional Repression in Monocytes: A Novel Mechanism of Communication between Inflammation and Metabolism. J. Immunol. 196, 2541–2551 (2016).

26. Mamas, M., Dunn, W. B., Neyses, L. & Goodacre, R. The role of metabolites and metabolomics in clinically applicable biomarkers of disease. Arch. Toxicol. 85, 5–17 (2011).

27. Rizzieri, D., Paul, B. & Kang, Y. Metabolic alterations and the potential for targeting metabolic pathways in the treatment of multiple myeloma. J. cancer metastasis Treat. 5, (2019).

28. Asgari, Y., Zabihinpour, Z., Salehzadeh-Yazdi, A., Schreiber, F. & Masoudi-Nejad, A. Alterations in cancer cell metabolism: The Warburg effect and metabolic adaptation. Genomics 105, 275–281 (2015).

29. Dufort, F. J. et al. Glucose-dependent de Novo Lipogenesis in B Lymphocytes. J. Biol. Chem. 289, 7011–7024 (2014).

30. Nakajima, K. et al. Glycolytic enzyme hexokinase II is a putative therapeutic target in B-cell malignant lymphoma. Exp. Hematol. 78, 46–55.e3 (2019).

31. Petrenko, O. et al. IL-6 promotes MYC-induced B cell lymphomagenesis independent of STAT3. PLoS One 16, e0247394 (2021).

32. Kim, J. H., Kim, W. S. & Park, C. Interleukin-6 mediates resistance to PI3K-pathway-targeted therapy in lymphoma. BMC Cancer 19, 1–11 (2019).

33. Stirm, K. et al. Tumor cell-derived IL-10 promotes cell-autonomous growth and immune escape in diffuse large B-cell lymphoma. Oncoimmunology 10, (2021).

34. Mariette, X. et al. Lymphoma in patients treated with anti-TNF: results of the 3-year prospective French RATIO registry. Ann. Rheum. Dis. 69, 400 (2010).

35. Cho, S. H. et al. Germinal centre hypoxia and regulation of antibody qualities by a hypoxia response system. Nature 537, 234–238 (2016).

36. Pera, B. et al. Metabolomic Profiling Reveals Cellular Reprogramming of B-Cell Lymphoma by a Lysine Deacetylase Inhibitor through the Choline Pathway. EBioMedicine 28, 80–89 (2018).

37. Arkatkar, T. et al. B cell-derived IL-6 initiates spontaneous germinal center formation during systemic autoimmunity. J. Exp. Med. 214, 3207–3217 (2017).

38. Bradshaw, P. C., Seeds, W. A., Miller, A. C., Mahajan, V. R. & Curtis, W. M. COVID-19: Proposing a Ketone-Based Metabolic Therapy as a Treatment to Blunt the Cytokine Storm. Oxid. Med. Cell. Longev. 2020, (2020).

39. Zheng, M. et al. The Cytokine Profiles and Immune Response Are Increased in COVID-19 Patients with Type 2 Diabetes Mellitus. J. Diabetes Res. 2021, (2021).

40. Lu, X. et al. Distinct IL-4-induced gene expression, proliferation, and intracellular signaling in germinal center B-cell-like and activated B-cell-like diffuse large-cell lymphomas. Blood 105, 2924–2932 (2005).

41. Zhou, L. et al. NEK2 Promotes Cell Proliferation and Glycolysis by Regulating PKM2 Abundance via Phosphorylation in Diffuse Large B-Cell Lymphoma. Front. Oncol. 11, 2071 (2021).

42. Chiche, J. et al. GAPDH Expression Predicts the Response to R-CHOP, the Tumor Metabolic Status, and the Response of DLBCL Patients to Metabolic Inhibitors. Cell Metab. 29, 1243–1257.e10 (2019).

43. Gupta, M. et al. Elevated serum IL-10 levels in diffuse large B-cell lymphoma: a mechanism of aberrant JAK2 activation. Blood 119, 2844–53 (2012).

44. Načinović-Duletić, A., Štifter, S., Dvornik, Š., Škunca, Ž. & Jonjić, N. Correlation of serum IL-6, IL-8 and IL-10 levels with clinicopathological features and prognosis in patients with diffuse large B-cell lymphoma. Int. J. Lab. Hematol. 30, 230–239 (2008).

45. Matsuda, T. & Kishimoto, T. Interleukin 6. Encycl. Immunol. 1458–1461 (1998). doi:10.1006/RWEI.1999.0371

46. Maeda, K., Mehta, H., Drevets, D. A. & Coggeshall, K. M. IL-6 increases B-cell IgG production in a feed-forward proinflammatory mechanism to skew hematopoiesis and elevate myeloid production. Blood 115, 4699 (2010).

47. Hashwah, H. et al. The IL-6 signaling complex is a critical driver, negative prognostic factor, and therapeutic target in diffuse large B-cell lymphoma. EMBO Mol. Med. 11, (2019).

48. Agrawal, S., Gollapudi, S., Su, H. & Gupta, S. Leptin activates human B cells to secrete TNF-α, IL-6, and IL-10 via JAK2/STAT3 and p38MAPK/ERK1/2 signaling pathway. J. Clin. Immunol. 31, 472–478 (2011).

49. Pauly, F. et al. Plasma immunoprofiling of patients with high-risk diffuse large B-cell lymphoma: a Nordic Lymphoma Group study. Blood Cancer J. 2016 611 6, e501–e501 (2016).

50. Li, R. et al. Cytokine-defined B cell responses as therapeutic targets in multiple sclerosis. Front. Immunol. 6, 626 (2016).

51. Rieckmann, P., D’Alessandro, F., Nordan, R. P., Fauci, A. S. & Kehrl, J. H. IL-6 and tumor necrosis factor-alpha. Autocrine and paracrine cytokines involved in B cell function. J. Immunol. 146, (1991).

52. Benihoud, K. et al. Respective roles of TNF-α and IL-6 in the immune response-elicited by adenovirus-mediated gene transfer in mice. Gene Ther. 2007 146 14, 533–544 (2006).

53. Mauer, J., Denson, J. L. & Brü, J. C. Versatile functions for IL-6 in metabolism and cancer. Trends Immunol. 36, 92–101 (2015).

54. Linge, I. et al. Pleiotropic Effect of IL-6 Produced by B-Lymphocytes During Early Phases of Adaptive Immune Responses Against TB Infection. Front. Immunol. 13, 137 (2022).

55. Schioppa, T. et al. B regulatory cells and the tumor-promoting actions of TNF-α during squamous carcinogenesis. Proc. Natl. Acad. Sci. U. S. A. 108, 10662–10667 (2011).

56. Zhang, D. et al. Metabolic regulation of gene expression by histone lactylation. Nature 574, 575–580 (2019).

57. Irizarry-Caro, R. A. et al. TLR signaling adapter BCAP regulates inflammatory to reparatory macrophage transition by promoting histone lactylation. Proc. Natl. Acad. Sci. U. S. A. 117, 30628–30638 (2020).

58. Chang, C. H. et al. XPosttranscriptional control of T cell effector function by aerobic glycolysis. Cell 153, 1239 (2013).

59. Galván-Peña, S. et al. Malonylation of GAPDH is an inflammatory signal in macrophages. Nat. Commun. 2019 101 10, 1–11 (2019).

60. Hitt, M. & Gauldie, J. Gene vectors for cytokine expression in vivo. Curr. Pharm. Des. 6, 613–632 (2000).

61. Vlasova-St. Louis, I. & Bohjanen, P. R. Post-Transcriptional Regulation of Cytokine Signaling by AU-Rich and GU-Rich Elements. J. Interf. Cytokine Res. 34, 233 (2014).

62. Chen, A. N. et al. Lactylation, a Novel Metabolic Reprogramming Code: Current Status and Prospects. Front. Immunol. 12, 2280 (2021).

63. Williams, N. C. & O’Neill, L. A. J. A role for the krebs cycle intermediate citrate in metabolic reprogramming in innate immunity and inflammation. Frontiers in Immunology 9, 1 (2018).

64. Zasłona, Z. & O’Neill, L. A. J. Cytokine-like Roles for Metabolites in Immunity. Mol. Cell 78, 814–823 (2020).

65. Martínez-Reyes, I. & Chandel, N. S. Mitochondrial TCA cycle metabolites control physiology and disease. Nat. Commun. 2020 111 11, 1–11 (2020).

66. Choi, I., Son, H. & Baek, J. H. Tricarboxylic Acid (TCA) Cycle Intermediates: Regulators of Immune Responses. Life 11, 1–19 (2021).

67. Chen, X., Song, M., Zhang, B. & Zhang, Y. Reactive Oxygen Species Regulate T Cell Immune Response in the Tumor Microenvironment. Oxid. Med. Cell. Longev. 2016, 1–10 (2016).

68. Murphy, M. P. How mitochondria produce reactive oxygen species. Biochem. J. 417, 1–13 (2009).

69. Patil, N. K., Bohannon, J. K., Hernandez, A., Patil, T. K. & Sherwood, E. R. Regulation of leukocyte function by citric acid cycle intermediates. J. Leukoc. Biol. 106, 105–117 (2019).

70. Hatzivassiliou, G. et al. ATP citrate lyase inhibition can suppress tumor cell growth. Cancer Cell 8, 311–321 (2005).

71. Dominguez, M., Brüne, B. & Namgaladze, D. Exploring the Role of ATP-Citrate Lyase in the Immune System. Front. Immunol. 12, 14 (2021).

72. Zhao, Y. et al. Citrate Promotes Excessive Lipid Biosynthesis and Senescence in Tumor Cells for Tumor Therapy. Adv. Sci. 9, 2101553 (2022).

73. Lu, X. et al. Distinct IL-4-induced gene expression, proliferation, and intracellular signaling in germinal center B-cell-like and activated B-cell-like diffuse large-cell lymphomas. Blood 105, 2924–2932 (2005).

74. Sarosiek, K. A., Nechushtan, H., Lu, X., Rosenblatt, J. D. & Lossos, I. S. Interleukin-4 Distinctively Modifies Responses of Germinal Center-like and Activated B-cell-like Diffuse Large B-cell Lymphomas to Immuno-chemotherapy. Br. J. Haematol. 147, 308 (2009).

75. Wallace, K. B., Sardão, V. A. & Oliveira, P. J. Mitochondrial Determinants of Doxorubicin-Induced Cardiomyopathy. Circ. Res. 126, 926–941 (2020).

