## Supplemental figure 1 and 2 for "Metabolic regulation of cytokine responses in diffuse large B-cell lymphoma"

**Figure s2.**

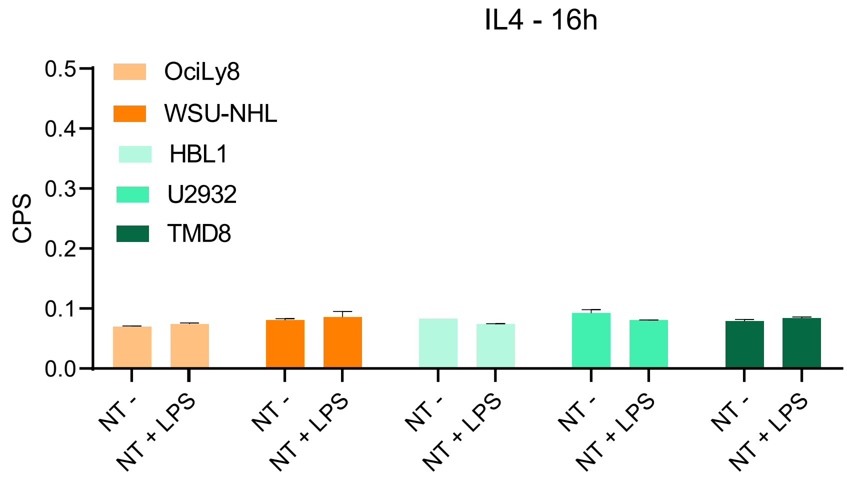
IL4 - 16h
CPS
0.5
0.4
0.3
0.2
0.1
0.0
OciLy8
WSU-NHL
HBL1
U2932
TMD8
NT -
NT + LPS
NT -
NT + LPS
NT -
NT + LPS
NT -
NT + LPS
NT -
NT + LPS

**Figure s1.**

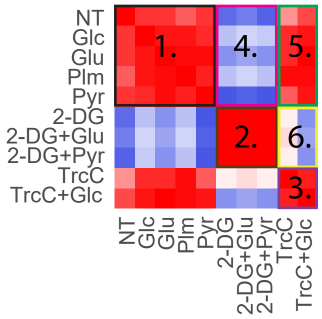


### Correlation Comparisons

1. Correlation between Metabolic Supplementations
2. Correlation between Glycolytic Inhibitions
3. Correlation between FA Inhibitions
4. Correlation between Metabolic Supplementation and Glycolytic Inhibition
5. Correlation between Metabolic Supplementation and FA inhibition
6. Correlation between Glycolytic Inhibition and FA inhibition

**1.** Metabolic Supplementations

| | | No LPS IL-6 | No LPS IL-10 | No LPS TNFa | With LPS IL-6 | With LPS IL-10 | With LPS TNFa |
|---|---|---|---|---|---|---|---|
| No LPS | IL-6 | 0.58 | 0.23 | 0.11 | 0.47 | 0.52 | 0.51 |
| | IL-10 | | 0.26 | -0.04 | 0.01 | 0.29 | 0.22 |
| | TNFa | | | 0.28 | 0.37 | -0.07 | 0.09 |
| With LPS | IL-6 | | | | 0.86 | 0.15 | 0.43 |
| | IL-10 | | | | | 0.67 | 0.44 |
| | TNFa | | | | | | 0.47 |

**2.** Glycolytic Inhibition

| | | No LPS IL-6 | No LPS IL-10 | No LPS TNFa | With LPS IL-6 | With LPS IL-10 | With LPS TNFa |
|---|---|---|---|---|---|---|---|
| No LPS | IL-6 | 0.92 | 0.85 | -0.44 | 0.87 | 0.83 | 0.42 |
| | IL-10 | | 0.97 | -0.61 | 0.79 | 0.88 | 0.32 |
| | TNFa | | | 0.78 | -0.31 | -0.33 | 0.24 |
| With LPS | IL-6 | | | | 0.99 | 0.80 | 0.57 |
| | IL-10 | | | | | 0.94 | 0.55 |
| | TNFa | | | | | | 0.80 |

**3.** FA Inhibition

| | | No LPS IL-6 | No LPS IL-10 | No LPS TNFa | With LPS IL-6 | With LPS IL-10 | With LPS TNFa |
|---|---|---|---|---|---|---|---|
| No LPS | IL-6 | 0.77 | 0.16 | 0.26 | 0.71 | -0.24 | 0.27 |
| | IL-10 | | 0.77 | 0.25 | 0.54 | 0.25 | -0.14 |
| | TNFa | | | 0.57 | 0.43 | -0.39 | 0.02 |
| With LPS | IL-6 | | | | 0.93 | -0.08 | 0.19 |
| | IL-10 | | | | | 0.82 | -0.12 |
| | TNFa | | | | | | 0.19 |

**4.** Metabolic Supplementations VS Glycolytic Inhibition

| | | No LPS IL-6 | No LPS IL-10 | No LPS TNFa | With LPS IL-6 | With LPS IL-10 | With LPS TNFa |
|---|---|---|---|---|---|---|---|
| No LPS | IL-6 | 0.13 | 0.11 | 0.29 | 0.28 | 0.32 | 0.64 |
| | IL-10 | | 0.19 | 0.12 | 0.39 | 0.29 | 0.35 |
| | TNFa | | | -0.44 | -0.30 | -0.31 | 0.05 |
| With LPS | IL-6 | | | | -0.41 | -0.26 | 0.32 |
| | IL-10 | | | | | 0.54 | 0.67 |
| | TNFa | | | | | | 0.56 |

**5.** Metabolic Supplementations VS FA Inhibition

| | | No LPS IL-6 | No LPS IL-10 | No LPS TNFa | With LPS IL-6 | With LPS IL-10 | With LPS TNFa |
|---|---|---|---|---|---|---|---|
| No LPS | IL-6 | 0.45 | 0.11 | 0.18 | 0.55 | 0.01 | 0.26 |
| | IL-10 | | 0.03 | 0.16 | 0.19 | 0.09 | 0.06 |
| | TNFa | | | 0.29 | 0.32 | -0.38 | 0.08 |
| With LPS | IL-6 | | | | 0.81 | -0.22 | 0.24 |
| | IL-10 | | | | | 0.19 | 0.19 |
| | TNFa | | | | | | 0.24 |

**6.** Glycolytic Inhibition VS FA Inhibition

| | | No LPS IL-6 | No LPS IL-10 | No LPS TNFa | With LPS IL-6 | With LPS IL-10 | With LPS TNFa |
|---|---|---|---|---|---|---|---|
| No LPS | IL-6 | -0.29 | -0.54 | -0.20 | -0.40 | 0.21 | 0.09 |
| | IL-10 | | -0.44 | -0.45 | -0.42 | 0.52 | 0.07 |
| | TNFa | | | 0.51 | 0.69 | 0.32 | 0.14 |
| With LPS | IL-6 | | | | -0.15 | -0.44 | 0.06 |
| | IL-10 | | | | | 0.35 | 0.20 |
| | TNFa | | | | | | 0.29 |
