## Supplemental text figure 1 ang 2 for "Metabolic regulation of cytokine responses in diffuse large B-cell lymphoma"

**Figure S1. Overview of average correlations between different metabolic treatments.**

Average correlations between the main metabolic treatment pathways and the effect they had on the different cytokines. For each cytokine, 6 different comparisons of interest could be made. Correlations indicate whether cytokine A and cytokine B responded similar to 1: metabolic supplementation, 2: glycolytic inhibition, 3: fatty acid (FA) inhibition, 4: metabolic supplementation versus glycolytic inhibition, 5: metabolic supplementation versus fatty acid inhibition and 6: glycolytic inhibition and fatty acid inhibition.

**Figure S2. Quantification of IL-4 production by GCB- and ABC-DLBCL cell lines.**

IL-4 production was measured in response to LPS for 2 GCB-DLBCL cell lines (orange) and three ABC-DLBCL cell lines (green). CPS (Counts per second).
